# Time-on-task-related decrements in performance in the rodent continuous performance test are not caused by physical disengagement from the task

**DOI:** 10.1101/2024.09.05.611415

**Authors:** Ye Li, Thomas van Kralingen, Megan Masi, Brandon Villanueva Sanchez, Beyonca Mitchell, Joshua Johnson, Jorge Miranda-Barrientos, Jason J. Rehg, Keri Martinowich, Gregory V. Carr

## Abstract

Attention deficits, a hallmark of many neuropsychiatric disorders, significantly impair quality of life and functional outcome for patients. Continuous Performance Tests (CPTs) are widely used to assess attentional function in clinical settings and have been adapted for mice as the rodent Continuous Performance Test (rCPT). In this study, we combined traditional analyses of rCPT performance with markerless pose estimation using DeepLabCut and visual field analysis (VFA) to objectively measure the orientation of mice toward stimuli during rCPT sessions. Additionally, we extended session lengths to assess performance decrements over time. Our findings show that extending rCPT sessions from 45 to 90 minutes results in a significant decline in performance in male mice, which aligns with performance decrements observed in clinical research. Importantly, physical engagement with the task remained relatively stable throughout the session, even as performance deteriorated. This suggests that the performance decline specifically reflects a time-on-task (TOT)-dependent vigilance decrement rather than physical disengagement. We also investigated the effects of amphetamine, an FDA-approved treatment for attention-deficit/hyperactivity disorder (ADHD), on rCPT performance. Amphetamine significantly improved rCPT performance in male mice by reducing false alarms without modulating orientation or physical engagement with the task stimuli. Collectively, these findings validate a behavioral tracking platform for objectively measuring physical engagement in the rCPT and a task modification that accentuates TOT-dependent performance decrements, enhancing the translational value of the rCPT for studies related to human neuropsychiatric disorders.

## INTRODUCTION

Attention is a fundamental cognitive domain that is disrupted in many neuropsychiatric disorders. The ability to sustain focus on specific tasks is essential for daily functioning and it is compromised in conditions such as attention deficit hyperactivity disorder (ADHD), schizophrenia [1–3], and mood disorders [4]. Understanding the underlying mechanisms of attention and its alterations in these disorders is imperative for developing effective therapeutic interventions.

Continuous Performance Tests (CPTs) were first designed to measure attention deficits in patients with traumatic brain injuries and have emerged as valuable tools for assessing attention function broadly across many neuropsychological settings [5–8]. In CPTs, the participants are required to respond to specific target stimuli while inhibiting responses to non-target stimuli. Because participants can produce both errors of commission (false alarms) and errors of omission (misses), data from CPTs can be analyzed using parameters derived from signal detection theory [9]. With the advent of computerized versions of CPTs, these tasks offer precise and accurate quantification of an individual’s attentional control, vigilance, and ability to process information under conditions requiring sustained concentration [8].

The development of touchscreen-based rodent CPTs (rCPT) has heralded new opportunities for exploring the brain mechanisms involved in attention and the impact of pharmacological interventions in rodent models [10,11]. Various adaptations of the rCPT have been employed to pinpoint the specific brain regions associated with attention function in mice and to examine the putative cognitive-enhancing effects of multiple drugs [11–16]. The utilization of the rCPT holds promise for increasing the translational utility of animal studies with respect to our understanding of attention function in humans.

In CPTs, Time-on-Task (TOT) is associated with a significant decline in performance [17]. Interestingly, this decline, referred to as a vigilance decrement, is more pronounced in patients with schizophrenia [18,19], suggesting that the TOT-induced vigilance decrement is a metric that may be useful for evaluating potential therapeutics for the treatment of attention deficits associated with schizophrenia. Although the exact biological mechanisms underlying TOT-induced vigilance decrements remain unclear, some evidence points to shifts in arousal, motivation, and mental fatigue, with mental fatigue understood as the exhaustion of information processing resources [20,21]. Therefore, further research into the biological foundations of TOT-related performance declines will require establishing and validating preclinical models. Moreover, confirming whether animal models capture the complex dynamics of human attention is particularly important for neuropsychiatric and pharmacological research purposes. Towards this goal, we set out to test whether we could induce a significant vigilance decrement by increasing the rCPT session length, which would allow us to provide a more in-depth investigation of TOT-dependent effects on performance.

The development of deep-learning-based behavioral analysis methodologies has enabled the markerless tracking of individual body parts in freely moving rodents with a precision that manual methods cannot achieve [22]. Utilization of neural network-based behavioral analysis tools, such as DeepLabCut (DLC)[23], allow for deep phenotyping beyond the traditional measures of task performance. A notable challenge with the rCPT is the difficulty in ascertaining the true engagement level of the animals during sessions. In contrast to humans, who can communicate their engagement and motivation levels to researchers, obtaining this information from rodents is difficult and can only be estimated indirectly. In this study, we describe a new platform that combines DLC with visual field analysis (VFA) [24] to objectively measure physical engagement with stimuli during rCPT sessions. Additionally, we test this platform by investigating the effect of amphetamine, an FDA-approved treatment for ADHD, on performance and physical engagement during TOT-rCPT sessions. We believe this novel approach has the potential to provide profound insights into the intricacies of rCPT performance in mice, enriching our understanding of attentional processes in a translational assay with utility in normal and pathological contexts.

## MATERIALS AND METHODS

### Mice

Male and female C57BL/6J mice (Strain # 000664; The Jackson Laboratory, Bar Harbor, ME, USA) were 9 weeks old at the start of the experiments. Mice were housed in polycarbonate cages (4 mice/cage) (Innovive, San Diego, CA, USA) in a room maintained on a 12h/12h light/dark cycle (lights on at 06:00 h). After handling mice three to five times (one time per day for one minute) over the course of a week, we started food (Teklad Irradiated Global 16% Protein Rodent Diet; #2916, Envigo, Indianapolis, IN, USA) restriction to maintain body weight between 85–90% of the average weight for C57BL/6J mice based on age and the growth curve provided by The Jackson Laboratory. Mice were provided with free access to drinking water throughout the experiments. We tested two separate cohorts of mice in this study. The first cohort consisted of six males and four females. The second cohort consisted of fifteen males and seven females. All procedures were approved by the Johns Hopkins University Animal Care and Use Committee and were in accordance with the *Guide for the Care and Use of Laboratory Animals*.

### Apparatus

All behavioral tasks were conducted in Bussey-Saksida touchscreen chambers (Model 80614E, Lafayette Instrument, Lafayette, IN, USA). Male and female mice were tested in separate chambers. To analyze pose tracking from video data, chambers were modified to optimize the quality of the video recordings (Figure 1). Briefly, the chamber cover was removed to provide an unobstructed view for the camera. Customized wall boosters were attached to the interior chamber to prevent mice from jumping out of the apparatus. The wall boosters were made of 0.56 cm thick transparent acrylic. The shape of the boosters matched the trapezoidal shape of the chamber and increased the overall height by 17 cm. To increase the video contrast, the default house lights in the chambers were replaced with led light strips attached on the top drawer and oriented toward the chamber. The camera location was moved to the center of the chamber.

**Figure 1.**
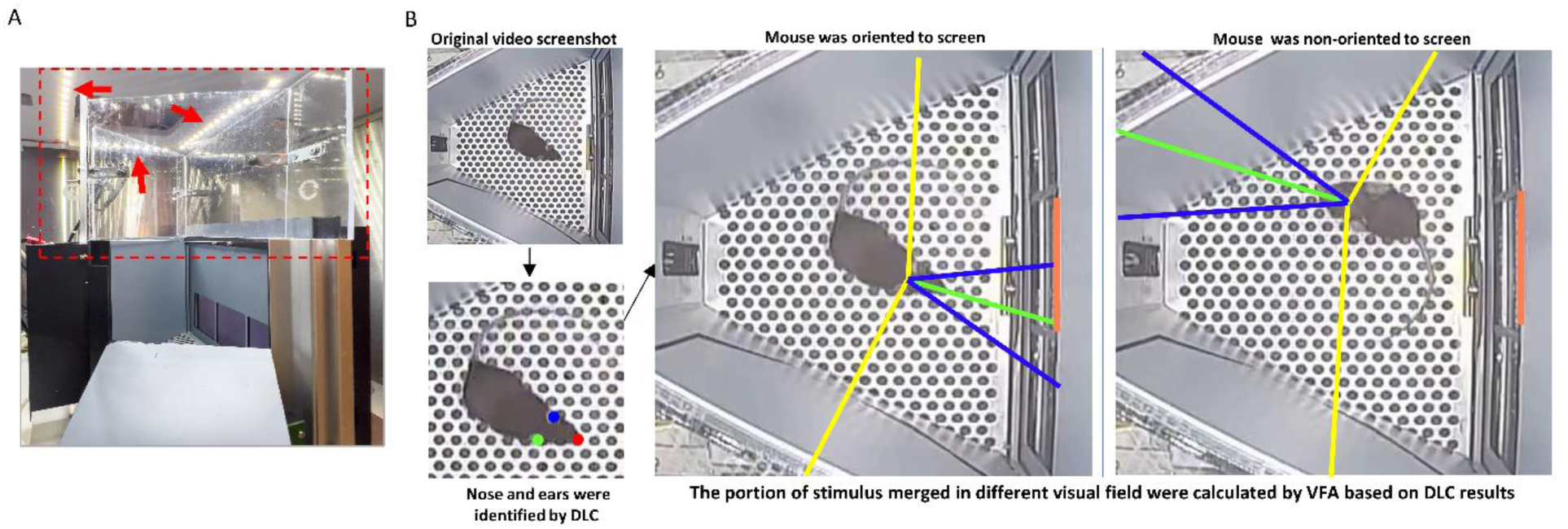
Customized touchscreen chamber and DLC-VFA analysis. **A** To increase the clarity of the video recording, the cover of the touchscreen chamber was replaced by a transparent wall booster (red dotted box). Additionally, the default house lights in the chambers were replaced with LED light strips, which were attached to the top drawer (red arrows). Additionally, the camera position was moved to the center of the chamber. **B** The top-down video was used for DLC and VFA analysis. In the illustrated video frame, DeepLabCut identified key anatomical points: the nose (marked with a red dot), the left ear (blue dot), and the right ear (green dot). Utilizing these points, the VFA package calculated the percentage of the stimulus area (the center hole area on the screen, marked with orange) that falls within each visual field area of the mouse. The green line is the center line of visual field. The area between blue lines is the binocular visual field. The areas between the blue line and yellow line are the monocular left or right visual fields. The screenshots show examples of frames when a mouse oriented and not oriented to the screen.

### The rCPT-training session

The rCPT protocol was based on previously published reports with some modifications [11,13–15; Figure 2]. Briefly, mice were exposed to strawberry milk (NESQUIK® Low Fat Strawberry Flavor Milk, Nestlé, Vevey, Switzerland) in their home cage for two days prior to introduction to the touchscreen chambers. For habituation sessions in the chambers, strawberry milk (200 µL) was loaded into the reward tray. A 3-window mask (Model 80614-M2) was placed in front of the touchscreen for all training and testing sessions. Rewards were not delivered when the screen was touched during habituation sessions. The criterion for advancement during habituation was consumption of all milk within 20 minutes over two consecutive sessions.

**Figure 2.**
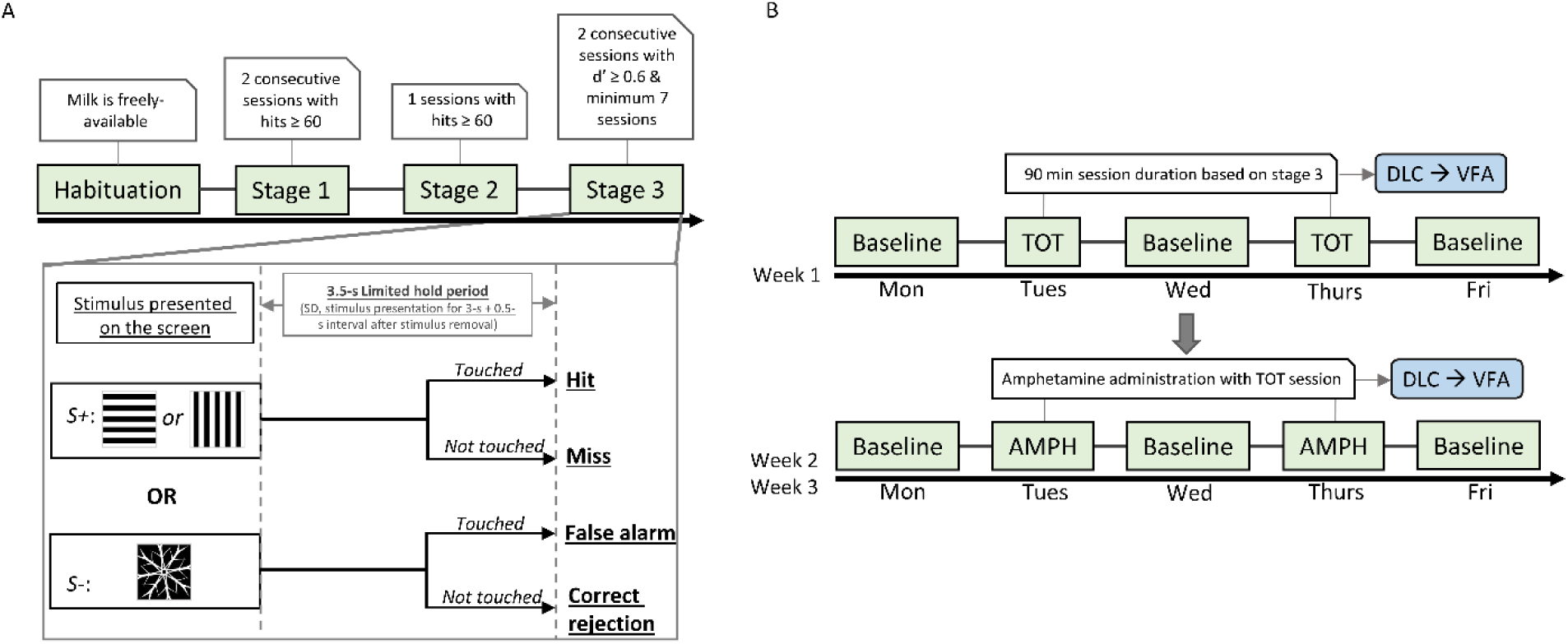
rCPT training stages with DeepLabCut (DLC) and VisualFieldAnalsysis (VFA) at Time-On-Task (TOT) and Amphetamine (AMPH) stages. **A** Timeline of rCPT training. Stages are denoted by green boxes and connected box indicates the required criterion of each stage to move mice to the next stage. In stage 3, both target (S+) or non-target (S-) were pseudorandomly presented with an average probability of 50% in session. The 3.5 second limited hold period started with S+ or S- presentation. Based on the response to the S+ or S-, four different events are possible: hit, miss, false alarm and correct rejection. **B** Timeline for TOT probe sessions and AMPH/vehicle treatment. Once mice reached Stage 3 criteria, they underwent a week of TOT probe testing followed by two weeks of amphetamine administration. Baseline tests occurred on Mondays, Wednesdays, and Fridays. TOT probes or AMPH/vehicle treatments were administered on Tuesdays and Thursdays. During the two-week amphetamine phase, four doses were tested (0 mg/kg/vehicle, 0.3 mg/kg, 0.6 mg/kg, and 1 mg/kg) following a Latin-square design. Additionally, TOT probe and amphetamine session videos were analyzed using DLC and VFA.

The formal rCPT training consists of three stages. In Stage 1, mice were trained to touch the center location during the limited hold period (LH), which was defined as the combination of the 10-second stimulus duration (SD) and a 0.5 second interval after the stimulus was removed from the screen. The stimulus was a white square image within a white-outlined frame (3.5×3.5 cm). A correct response toward this white square image triggered delivery of 20 µL of strawberry milk to the reward trough, the presentation of a 1-second 3 kHz tone, and illumination of the reward trough light. A head entry into the reward trough to collect the reward initiated an inter-trial interval (ITI) of 2s. Following the ITI, the next trial began with the presentation of the white square. The ITI would restart if the center location was touched during the ITI period. Mice were required to earn 60 rewards in a 45-minute session to advance to Stage 2.

In Stage 2, the white square image was replaced with the S+ stimulus (target), which was either an array of horizontal or vertical alternating black and white bars. The S+ image was the same size as the white square image from Stage 1. Mice were pseudorandomly assigned to one of the two S+ images to balance within sex. Also in Stage 2, the SD was reduced to 5 s. All other parameters and advancement criteria were the same as Stage 1.

In stage 3, a novel S- (snowflake image, non-target) stimulus was introduced to mice. Both S+ and S- mice were pseudorandomly presented with an average probability of 50%. The SD period was reduced to 3s. A response to the S+ during the LH period was recorded as a “hit”. If mice did not respond to the S+ during the LH, a “miss” was recorded. If mice touched the screen during the LH when the S- was presented, a “false alarm” was recorded. If mice did not touch the screen when the S- was presented, a “correct rejection” was recorded. False alarms were followed by correction trials. In correction trials, the S- stimulus was always presented. The mouse was required to withhold responding to the S-. If the mouse responded, then the ITI would reset and another correction trial would commence. Mice would stay within the correction trial loop until they withheld responding. All of the other response types (Hits, Misses, and Correct rejection) were followed by an ITI period. The ITI in stage 3 was increased to five seconds. To pass this stage, mice needed to discriminate between S+ and S-, with a d’ > 0.6 for two consecutive sessions and complete a minimum of seven total sessions (see data analysis section for definition of d’). Mice were trained Monday-Friday with no training or testing on Saturday or Sunday. If mice reached criterion on days other than Friday, they were kept on Stage 3 training until the following Monday.

### Time-On-Task Probe

The design of Time-On-Task (TOT) probe sessions was based on stage 3 of the rCPT with a few modifications. First, the duration of the session was increased from 45 minutes to 90 minutes. Next, the maximum reward number was changed from 150 to 999, to ensure mice were not able to reach the cap during a session. After completing Stage 3 training, mice started the TOT probe test phase. The TOT probe tests were conducted on Tuesday and Thursday. On Monday, Wednesday and Friday, performance was tested in baseline (45 minutes) rCPT sessions, with parameters identical to Stage 3

### Drug administration

Amphetamine hemisulfate (AMPH) (Product number: A5880, Sigma-Aldrich, St. Louis, MO, USA) was dissolved in 0.9% saline and injected (i.p.; 10 ml/kg injection volume) 15 minutes before behavioral testing. Solutions were prepared fresh on each testing day. AMPH doses used in this study were 0.3, 0.6, and 1 mg/kg based on the free base. Doses were chosen based on previous studies from other groups and in-house pilot studies [13,14]. Each mouse received each dose of AMPH during testing. Dose order was determined according to a Latin-square design.

### Time-On-Task probe during Amphetamine study

AMPH was tested in TOT probe sessions. Drug testing sessions, including saline control sessions, were run on Tuesday and Thursday while sessions on Monday, Wednesday, and Friday were baseline Stage 3 sessions without any injection.

### Experimental procedure

Cohort 1 and Cohort 2 were tested using similar experimental procedures with a few differences. First, both cohorts advanced through rCPT-training and baseline testing sessions. In Cohort 1, once mice reached the Stage 3 performance criteria, they were tested on TOT and degraded stimulus (DS) probe sessions. The group was split so that half started with TOT and the other half started with DS. The DS data is included as the replication cohort in a separate publication [15]. Cohort 1 testing was complete at the end of the TOT and DS probe sessions. Cohort 2 were not testing in DS probe sessions and following their TOT probe sessions, they entered the drug-testing cycle with AMPH.

### DeepLabCut and Visual field analysis

Briefly, the rCPT videos were initially preprocessed and subsequently analyzed using the (DLC) package, version 2.2 [23,25] to estimate the positions of specific body parts, including the nose, left side of the head, and right side of the head. Based on the results derived from the DLC pipeline, further analysis was conducted using VFA [24]. Next, an orientation index was calculated to estimate the animal’s orientation relative to the stimulus displayed on the screen. Additional details are provided in the Supplemental Information section.

### Data analysis

All rCPT behavioral data were exported as .csv files from the ABET II application (Lafayette Instruments, Lafayette, IN, USA) prior to processing in Jupyter Notebook with Python (Version 3.7.6). Key metrics such as hit rate (HR), false alarm rate (FAR), sensitivity (d’), and response criterion (C) were calculated. For detailed parameter definitions and calculations, refer to Box 1.

#### Box 1

Parameters in rCPT and performing calculations based on signal detection theory principles.

**Hit:** correct touch to the S+ (target).

**Miss:** omission to the S+ (target).

**False alarm:** incorrect touch to the S- (non-target).

**Correct rejection:** to withhold response when a S- (non-target) is presented.

**Hit rate (HR)** = Hits/(Hits+Misses)

**False alarm rate (FAR)** = False alarms/(False alarms + Correct rejections)

**Sensitivity (d’)** = z(Hit rate)-z(False alarm rate)

**Response criterion (c)** = -(z(Hit rate)+z(False alarm rate))/2

**Impulsivity %** = (Centre ITI touches/Total ITIs initiated) x 100

**Response rate =** (Hits + False alarms)/(Hits + False alarms + Misses + Correct rejections)

The following packages were imported into a Jupyter Notebook on the Windows 10 platform: pandas (Version 1.0.1), numpy (Version 1.18.1), matplotlib (Version 3.1.3), seaborn (Version 0.11.0), scipy (Version 1.4.1). A linear mixed effect model was used to analyze the data by lmer function from *lme4* library (Version 1.1.35.1) in Rstuido (RStudio 2023.12.1+402, R version 4.3.3). The parameters (such as d’, c, HR, FAR et. al) were analyzed by the related main effects and the random effect of mouse ID. For example, in stage 3 training, the following model was used:

*Stage 3: parameter ∼ Session order + Sex + Session order * Sex + (1| Mouse ID).* Post hoc analyses were conducted using Bonferroni’s test, where appropriate. The significance threshold was set at p < 0.05 for all statistical analyses. For data visualization, we used the matplotlib-based seaborn package in Jupyter Notebook. All code used for data analysis is available on <u>GitHub (</u>https://github.com/yestmd/CPT-DLC-VFA).

## RESULTS

### Males and females reach criteria in the rCPT at the same rate

Both male and female mice improve their performance, as measured by d’ across Stage 3 training sessions (F(6,180) = 54.5160, *p* < 0.0001) and there were no sex differences in performance (F(1, 30) = 0.0330, *p* = 0.8570; Figure 3). Post hoc analyses indicated that d’ reached a plateau starting at the fourth training session (no significant differences between sessions 4-7 (Figure 3). The increase in d’ across training was driven by both a decrease in the FAR (F(6,180) = 18.0549, *p* < 0.0001) (Figure 3) and an increase in the hit rate (HR) (F(6, 180) = 10.2625, *p* = 0.0001; Figure 3).

**Figure 3.**
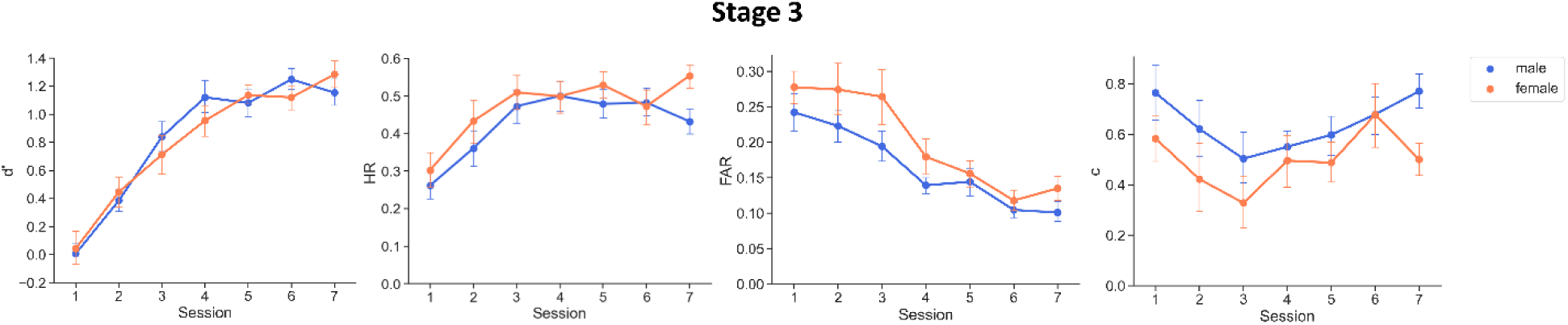
No sex differences in performance during Stage 3 training. Both male and female mice improve performance as measured by d’ across Stage 3 training sessions. The HR increases and the FAR decreases across sessions. Data are shown as mean ± SEM. n = 21 males and 11 females.

Interestingly, there was a main effect of training on the response bias metric c (F(6,180) = 2.3010, *p* = 0.0364; Figure 3). However, further post hoc analysis did not reveal any significant differences between individual training sessions. The data indicate that the response bias becomes more conservative across training. Additionally, there was no significant sex difference in c (F(1, 30) = 2.5399, *p* = 0.1215) (Figure 3).

Both correct (hits) and incorrect (false alarms) response latencies decreased across training as mice became more proficient in the task (F(6, 180) = 4.1646, *p* = 0.0006 and F(6, 180) = 2.5643, *p* = 0.0208, respectively). Interestingly, female mice responded faster than males on both hits (F(1, 30) = 7.0915, *p* = 0.0123) and false alarms (F(1, 30) = 4.9101, *p* = 0.0344; Table 1). In contrast, there were no significant differences in the reward latency, the amount of time between screen touch and reward retrieval, across either training or sex (F(6, 180) = 0.5663, *p* = 0.7568; F(1,30) = 0.0045, *p* = 0.9468, respectively). Impulsivity, measured as premature screen touches during ITIs, decreased over training (F(6, 180) = 3.5579, *p* = 0.0024; Supplemental Figure 1).

**Table 1.**
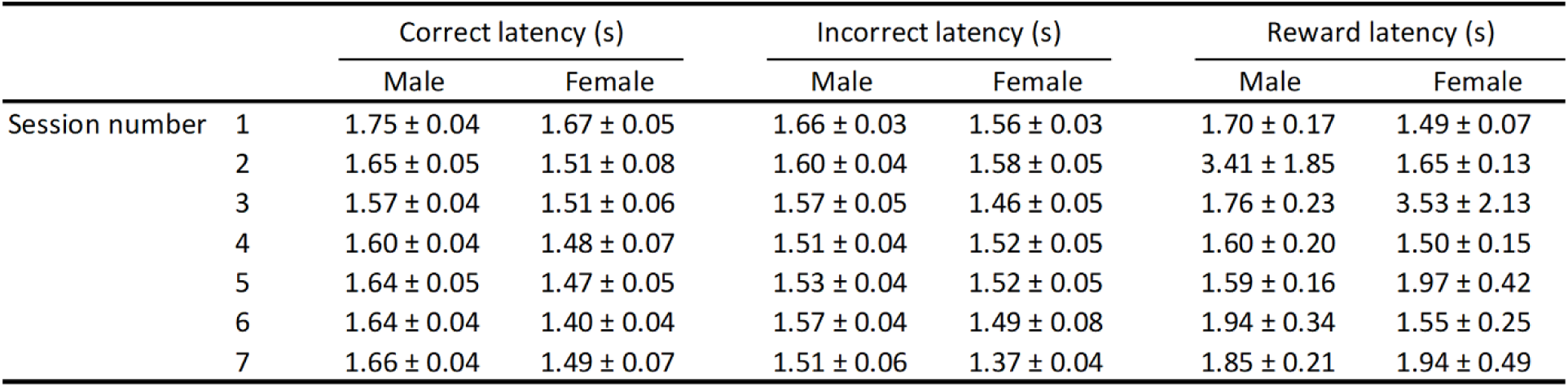

|  |  | Correct latency (s) |  | Incorrect latency (s) |  | Reward latency (s) |  |
| --- | --- | --- | --- | --- | --- | --- | --- |
|  |  | Male | Female | Male | Female | Male | Female |
| Session number | 1 | 1.75 ± 0.04 | 1.67 ± 0.05 | 1.66 ± 0.03 | 1.56 ± 0.03 | 1.70 ± 0.17 | 1.49 ± 0.07 |
|  | 2 | 1.65 ± 0.05 | 1.51 ± 0.08 | 1.60 ± 0.04 | 1.58 ± 0.05 | 3.41 ± 1.85 | 1.65 ± 0.13 |
|  | 3 | 1.57 ± 0.04 | 1.51 ± 0.06 | 1.57 ± 0.05 | 1.46 ± 0.05 | 1.76 ± 0.23 | 3.53 ± 2.13 |
|  | 4 | 1.60 ± 0.04 | 1.48 ± 0.07 | 1.51 ± 0.04 | 1.52 ± 0.05 | 1.60 ± 0.20 | 1.50 ± 0.15 |
|  | 5 | 1.64 ± 0.05 | 1.47 ± 0.05 | 1.53 ± 0.04 | 1.52 ± 0.05 | 1.59 ± 0.16 | 1.97 ± 0.42 |
|  | 6 | 1.64 ± 0.04 | 1.40 ± 0.04 | 1.57 ± 0.04 | 1.49 ± 0.08 | 1.94 ± 0.34 | 1.55 ± 0.25 |
|  | 7 | 1.66 ± 0.04 | 1.49 ± 0.07 | 1.51 ± 0.06 | 1.37 ± 0.04 | 1.85 ± 0.21 | 1.94 ± 0.49 |

### TOT affected rCPT performance in both males and females

To assess the effect of TOT on performance, mice were given 90-min probe sessions and these sessions were divided into six 15-min time bins for analysis (Figure 4). There were significant effects of time bin and sex on d’ (F(5, 150) = 8.0595, *p* <0.0001; F(1, 30) = 6.7372, *p* = 0.0145, respectively) and a significant time X sex interaction (F(5, 150) = 3.7817, *p* = 0.0030). Male mice showed a larger TOT-dependent decrement in performance (d’) than female mice and this was statistically significant in the last three time bins (bin4, male vs female, *p* = 0.0033; bin5, male vs female, *p* = 0.0001; bin6, male vs female, *p* = 0.0250). This difference appears to be due to a larger decrease in HR in male mice across time (HR: main effect of time bin, F(5, 150) = 45.2060, *p* < 0.0001; main effect of sex, F(1, 30) = 14.9250, *p* = 0.0006; time bin X sex, F(5, 150) = 2.2760, *p* = 0.0499). Additionally, response biases became more conservative across time in both males and females (main effect of time bin, F(5, 150) = 46.7010, *p* < 0.0001), although females had more liberal response biases across all time bins (main effect of sex, F(1, 30) = 18.1640, p= 0.0002; Figure 4).

**Figure 4.**
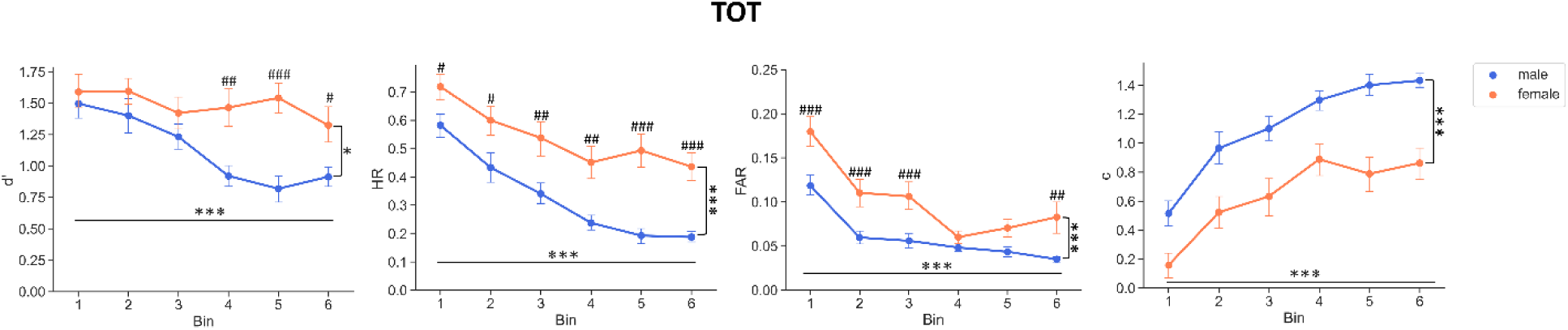
rCPT performance was affected by TOT. TOT probe sessions were divided into six equal time bins of 15 minutes. Performance decreased across time and there was a significant time bin X sex effect. Data are shown as mean ± SEM. n = 21 males and 11 females. An asterisk (*) indicates significant effects of time or sex. A hash (#) indicates significant pos hoc results of time bin X sex when significant interaction shown. \**p*<0.05, \*\*\**p*<0.001, ^#^*p*<0.05, ^##^*p*<0.01, ^###^*p*<0.001.

Correct response latency decreased over time while there was no change in the incorrect response latency (F(5, 150) = 3.2820, *p* = 0.0077; F(5, 150) = 0.4350, *p* = 0.8236, respectively; Table 2). Reward collection latency also increased slightly over time F(5, 150) = 2.3345, *p* = 0.0448; Table 2). Sex had no effect on correct response latency, incorrect response latency, or reward latency (F(1, 30) = 3.3690, *p* = 0.0764; F(1, 30) = 0.3496, p = 0.5588; F(1, 30) = 0.3817, *p* = 0.54137, respectively; Table 2). Moreover, premature responses decreased across time bins (main effect of time bin, F(5, 150) = 22.5124, *p* < 0.0001; Supplemental Figure 2) with a significant main effect of sex (F(1, 30) = 13.4867, *p* = 0.0009; Supplemental Figure 2). There was also a significant time bin X sex interaction effect, indicating that female mice had more premature responses in the first three time bins and also in bin 5 (F(5, 150) = 2.4092, *p* = 0.0391, post hoc male vs female at bin 1, *p* < 0.0001; bin 2, *p* = 0.0003; bin 3, *p* = 0.0060; bin 5, *p* = 0.0297; Supplemental Figure 2).

**Table 2.**

|  |  | Correct latency (s) |  | Incorrect latency (s) |  | Reward latency (s) |  |
| --- | --- | --- | --- | --- | --- | --- | --- |
|  |  | Male | Female | Male | Female | Male | Female |
| Time bin | 1 | 1.46 ± 0.06 | 1.30 ± 0.08 | 1.44 ± 0.06 | 1.29 ± 0.07 | 1.24 ± 0.07 | 1.06 ± 0.05 |
|  | 2 | 1.51 ± 0.06 | 1.40 ± 0.07 | 1.11 ± 0.08 | 1.41 ± 0.09 | 2.03 ± 0.69 | 1.26 ± 0.16 |
|  | 3 | 1.52 ± 0.05 | 1.46 ± 0.09 | 1.46 ± 0.13 | 1.24 ± 0.07 | 7.92 ± 5.67 | 1.37 ± 0.12 |
|  | 4 | 1.67 ± 0.08 | 1.50 ± 0.09 | 1.35 ± 0.11 | 1.47 ± 0.13 | 5.23 ± 1.75 | 5.86 ± 4.47 |
|  | 5 | 1.65 ± 0.08 | 1.47 ± 0.09 | 1.45 ± 0.12 | 1.29 ± 0.14 | 12.37 ± 7.14 | 7.53 ± 4.40 |
|  | 6 | 1.66 ± 0.07 | 1.52 ± 0.05 | 1.36 ± 0.13 | 1.22 ± 0.10 | 13.05 ± 6.61 | 5.75 ± 2.80 |

### AMPH attenuates the TOT-dependent decrease in performance and produces a more conservative response bias

In male mice, AMPH significantly increased d’ (F(3, 293.39) = 2.9648, *p* = 0.0324), with the 0.6 mg/kg dose improving performance compared to vehicle treatment (*p* = 0.0210). The improvement in performance in male mice was most likely due to a significant decrease in the FAR (F(3, 294.63) = 3.2999, *p* = 0.0208), with the effect of the 0.3 mg/kg dose being significantly different from vehicle (*p*= 0.0338). There were no effects of AMPH on HR or c (F(3, 293.20) = 0.6034, *p* = 0.6132; F(3, 293.41) = 0.8102, *p* = 0.4890), respectively) (Figure 5A).

**Figure 5.**
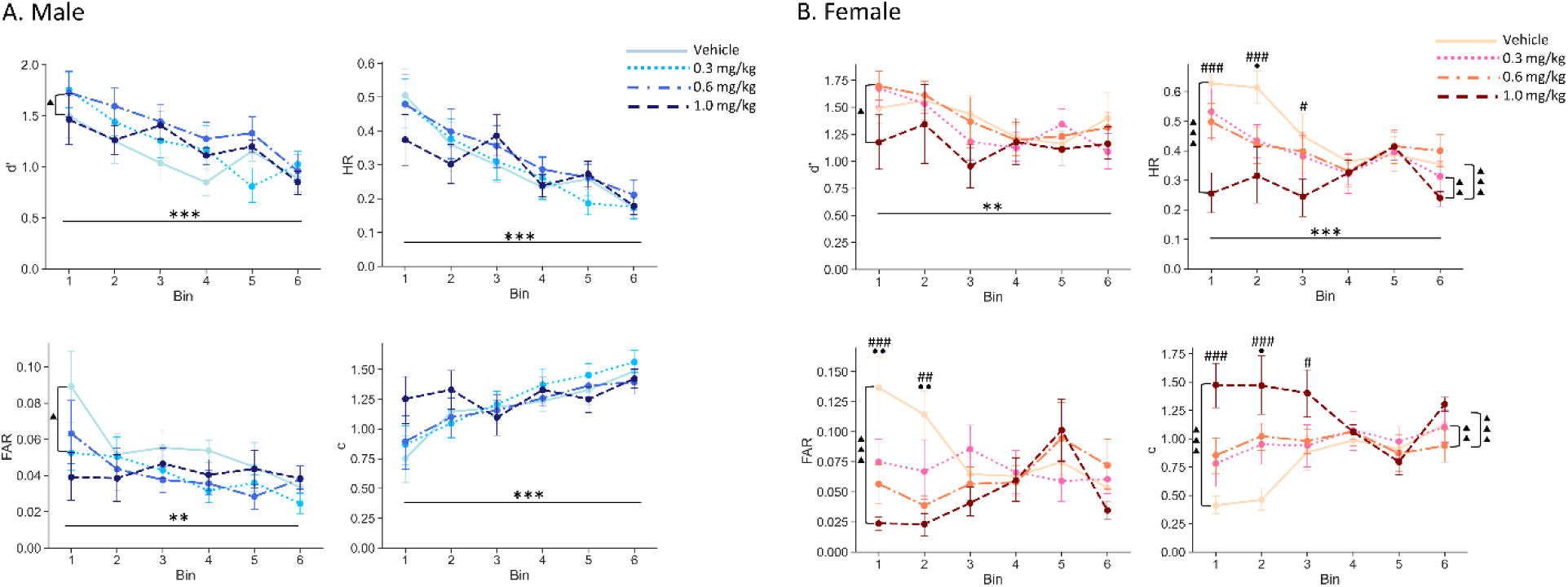
Effects of AMPH on rCPT performance. **A** AMPH at 0.6 mg/kg attenuates the TOT-dependent decrease in d’ in male mice. A triangle (▴) indicates significant pos hoc results between 0.6mg/kg and vehicle. An asterisk (*) indicates statistical significance of main effect of time bin or dosage. **B** Female mice showed significant increased C, decrease HR and FAR in highest dosage (1mg/kg). Further post hoc analysis also shown these changes mainly happened at first three time bin. Data are shown as mean ± SEM. n = 15 males and 7 females. An asterisk (*) indicates statistical significance of main effect of time bin or dosage. A hash (#) indicates significant effect of 1mg/kg comparing to vehicle. A circle (•) indicates a significant effect of 0.6mg/kg comparing to vehicle. \**p*<0.05, \*\*\**p*<0.001, ^#^*p*<0.05, ^##^*p*<0.01, ^###^*p*<0.001, • *p*<0.05, •• *p*<0.01.

In contrast, in female mice, AMPH decreased both the HR and FAR (F(3, 137.01) = 11.9560, *p* < 0.0001; F(3, 137.02) = 5.2299, *p* = 0.0019, respectively). Post hoc analyses indicated that the 1 mg/kg dose significantly decreased HR and FAR compared to vehicle treatment (p<0.0001, p=0.0009, respectively) and decreased the HR when compared to the 0.3 mg/kg (p=0.004) and 0.6 mg/kg (p=0.0008) doses. There were also significant time X treatment interactions for both HR and FAR (F(15, 137) = 2.0239, *p* = 0.0176; F(15, 137.02)=2.5240, p= 0.0025, respectively). Post hoc analyses showed that the both 0.6 mg/kg and 1mg/kg dose decreased the FAR in Bins 1 and 2 (0.6mg/kg: Bin1 p= 0.0042; Bin2 p= 0.0082; 1mg/kg: Bin1 p<0.0001; Bin2 p=0.0016), while 1mg/kg dose decreased the HR in Bin 1, Bin 2 and Bin 3 (p<0.0001; p= 0.0002; p= 0.0227, respectively) and 0.6 mg/kg decreased the HR in Bin 2 (p= 0.0390). Although, in female mice, there was a main effect of AMPH on d’ (F(3, 137.01)=3.5081, p=0.0171), post hoc analysis showed there was only a significant difference between the 0.6mg/kg and 1mg/kg doses (*p* = 0.0232) and no differences between any drug doses and vehicle treatment. On the other hand, AMPH also increased c in female mice (F(3, 137.01) = 12.7117, *p* < 0.0001). Post hoc analyses indicated that 1 mg/kg significantly increased c compared to vehicle treatment (*p* < 0.0001), 0.3 mg/kg *p* = 0.0015) and 0.6 mg/kg (*p* = 0.0007). A time × treatment interaction was identified (F(15, 137.01) = 2.7991, *p* = 0.0008). Compared to vehicle treatment, 1 mg/kg significantly increased c during Bin 1, Bin 2, and Bin 3 (*p* < 0.0001, *p* < 0.0001, *p* = 0.0305, respectively), while 0.6 mg/kg increased c at Bin 2 (*p* = 0.0160). Additionally, significant differences were found between 0.3 mg/kg and 1 mg/kg during Bin 1 and Bin 2 (*p* = 0.0014, *p* = 0.0365, respectively) and between 0.6 mg/kg and 1 mg/kg, during Bin 1 (p=0.0055).(Figure 5B).

AMPH treatment did not alter correct response latency, incorrect response latency, or reward latency in male mice (F(3, 295.32) = 1.2934, *p* = 0.2768; F(3, 306) = 1.5440, *p* = 0.2031; F(3, 295.28) = 0.7941, *p* = 0.4980, respectively; Table 3). We also analyzed response time standard deviations (SDs) as a measure of reaction time variability and there was a significant interaction effect between time bins and AMPH treatment on the incorrect latency SD in male mice (F(15, 291.61) = 1.7101, *p* = 0.0483). The 1 mg/kg dose reduced the incorrect latency SD during Bins 1 and 2 (*p* = 0.0190; *p* = 0.0491, respectively) (Supplemental Table 3). Similar to males, there was no significant effect of treatment on correct latency, incorrect latency and reward latency in female mice (F(3, 138) = 0.4190, *p* = 0.7397; F(3, 138) = 1.6233, *p* = 0.1868; F(3, 138) = 1.1092, *p* = 0.3476, respectively), but there was a significant main effect of treatment on incorrect latency SD (F(3, 138) = 3.1595, *p* = 0.0268), with 1 mg/kg dose decreasing the incorrect latency SD (*p* = 0.0155), just as it did in male mice.

**Table 3.**

| Male |  |  |  |  |  |  | Female |  |  |  |  |  |  |
| --- | --- | --- | --- | --- | --- | --- | --- | --- | --- | --- | --- | --- | --- |
| Dosage (mg/kg) | bin1 | bin2 | bin3 | bin4 | bin5 | bin6 | Dosage (mg/kg) | bin1 | bin2 | bin3 | bin4 | bin5 | bin6 |
|  | Correct latency (s)* |  |  |  |  |  |  | Correct latency (s) |  |  |  |  |  |
| 0 | 1.43 ± 0.14 | 1.49 ± 0.14 | 1.65 ± 0.07 | 1.70 ± 0.08 | 1.69 ± 0.09 | 1.72 ± 0.09 | 0 | 1.43 ± 0.11 | 1.38 ± 0.09 | 1.45 ± 0.04 | 1.50 ± 0.07 | 1.51 ± 0.06 | 1.67 ± 0.16 |
| 0.3 | 1.60 ± 0.08 | 1.71 ± 0.07 | 1.68 ± 0.06 | 1.68 ± 0.07 | 1.72 ± 0.12 | 1.62 ± 0.10 | 0.3 | 1.46 ± 0.08 | 1.48 ± 0.11 | 1.39 ± 0.09 | 1.60 ± 0.07 | 1.61 ± 0.10 | 1.69 ± 0.10 |
| 0.6 | 1.58 ± 0.12 | 1.53 ± 0.08 | 1.59 ± 0.10 | 1.57 ± 0.08 | 1.72 ± 0.11 | 1.57 ± 0.08 | 0.6 | 1.48 ± 0.09 | 1.51 ± 0.10 | 1.62 ± 0.10 | 1.44 ± 0.07 | 1.63 ± 0.10 | 1.63 ± 0.06 |
| 1 | 1.45 ± 0.19 | 1.55 ± 0.14 | 1.77 ± 0.09 | 1.63 ± 0.08 | 1.77 ± 0.07 | 1.65 ± 0.06 | 1 | 1.37 ± 0.27 | 1.23 ± 0.33 | 1.60 ± 0.34 | 1.63 ± 0.10 | 1.56 ± 0.07 | 1.49 ± 0.06 |
|  | Incorrect latency (s)** |  |  |  |  |  |  | Incorrect latency (s) |  |  |  |  |  |
| 0 | 1.15 ± 0.15 | 1.79 ± 0.20 | 1.44 ± 0.18 | 1.67 ± 0.15 | 1.09 ± 0.19 | 1.27 ± 0.21 | 0 | 1.09 ± 0.16 | 1.41 ± 0.06 | 1.18 ± 0.21 | 1.39 ± 0.15 | 1.48 ± 0.19 | 1.34 ± 0.40 |
| 0.3 | 1.04 ± 0.18 | 1.30 ± 0.22 | 1.25 ± 0.25 | 1.23 ± 0.25 | 1.32 ± 0.16 | 1.21 ± 0.24 | 0.3 | 0.76 ± 0.22 | 1.04 ± 0.25 | 0.91 ± 0.24 | 1.23 ± 0.27 | 1.48 ± 0.19 | 1.08 ± 0.22 |
| 0.6 | 0.88 ± 0.18 | 1.09 ± 0.25 | 1.27 ± 0.24 | 1.13 ± 0.17 | 1.05 ± 0.27 | 1.50 ± 0.18 | 0.6 | 0.97 ± 0.25 | 1.28 ± 0.27 | 1.41 ± 0.28 | 1.18 ± 0.13 | 1.14 ± 0.20 | 1.31 ± 0.23 |
| 1 | 0.65 ± 0.16 | 1.08 ± 0.30 | 1.27 ± 0.23 | 1.21 ± 0.23 | 1.10 ± 0.18 | 1.98 ± 0.17 | 1 | 0.62 ± 0.27 | 0.84 ± 0.46 | 0.83 ± 0.25 | 1.04 ± 0.30 | 1.36 ± 0.18 | 1.45 ± 0.38 |
|  | Reward latency (s) |  |  |  |  |  |  | Reward latency (s) |  |  |  |  |  |
| 0 | 1.06 ± 0.11 | 1.15 ± 0.13 | 1.68 ± 0.26 | 1.78 ± 0.23 | 1.76 ± 0.18 | 1.72 ± 0.18 | 0 | 1.09 ± 0.09 | 1.36 ± 0.14 | 2.12 ± 0.83 | 1.89 ± 0.40 | 1.95 ± 0.55 | 2.85 ± 1.11 |
| 0.3 | 1.59 ± 0.31 | 1.48 ± 0.14 | 1.85 ± 0.30 | 1.56 ± 0.18 | 1.83 ± 0.29 | 1.67 ± 0.15 | 0.3 | 1.20 ± 0.07 | 1.40 ± 0.10 | 1.77 ± 0.40 | 1.65 ± 0.22 | 3.38 ± 2.20 | 4.24 ± 2.25 |
| 0.6 | 1.63 ± 0.41 | 2.27 ± 1.00 | 1.83 ± 0.51 | 1.55 ± 0.20 | 1.77 ± 0.28 | 1.44 ± 0.16 | 0.6 | 1.45 ± 0.11 | 1.98 ± 0.45 | 4.12 ± 2.56 | 4.21 ± 2.88 | 1.45 ± 0.27 | 2.33 ± 1.14 |
| 1 | 1.43 ± 0.28 | 2.17 ± 0.74 | 2.14 ± 0.53 | 1.59 ± 0.18 | 1.62 ± 0.14 | 1.77 ± 0.20 | 1 | 0.94 ± 0.25 | 1.25 ± 0.39 | 264.24 ± 254.33 | 5.38 ± 3.71 | 5.09 ± 2.11 | 4.06 ± 1.43 |

AMPH decreased premature responding in male mice (Time X treatment interaction effect; (F(15, 292.19) = 2.0621, *p* = 0.0118) with the 0.6 and 1 mg/kg doses decreasing premature responses in Bin 1 (*p* = 0.008 and *p* < 0.0001, respectively) compared to vehicle treatment. In female mice, there was also a significant time X treatment interaction (F(15, 137.01) = 2.4443, *p* = 0.0034), with the 0.6 and 1 mg/kg doses decreasing impulsivity % during Bin 1 (*p* = 0.0022; *p* < 0.0001, respectively) and Bin 2 (*p* = 0.0429; *p* = 0.0148, respectively) compared to vehicle treatment (Supplemental Figure 3).

### The TOT-dependent performance decrement is not due to changes in orientation to the touchscreen

We next tested whether the effects of TOT were due to changes in orientation toward the touchscreen during trials. In other words, could the decrease in d’ and response rate be due to mice engaging in activities unrelated to touchscreen stimulus presentation (e.g. exploring the chamber or grooming). We developed an orientation index to provide quantitative data related to time spent oriented toward the touchscreen. Also, to measure active engagement with the touchscreen stimuli, we introduced a metric termed “response rate”. The response rate is a composite of the HR and FAR (i.e. active responses), calculated by dividing the total number of hits and false alarms by the total number of all events (hits, false alarm, misses and correct rejections). The total orientation index of male mice was relatively stable (∼0.9) across sessions. There was a small, but statistically significant decrease in orientation across time (F(5, 100) = 3.6555, *p* = 0.0044). However, the only significant difference was between Bin 1 and Bin 6 (*p*=0.0016; Figure 6A). In contrast, the response rate began to decline almost immediately (F(5, 100) = 47.1140, *p* < 0.0001; post hoc analyses: Bin1 vs Bin2, *p* < 0.0001; Bin1 vs Bin3, *p* < 0.0001; Bin1 vs Bin4, *p* < 0.0001; Bin1 vs Bin5, *p* < 0.0001; Bin1 vs Bin6, *p* < 0.0001). This finding suggests the decrease in response rate is not explained by a concomitant decrease in orientation.

**Figure 6.**
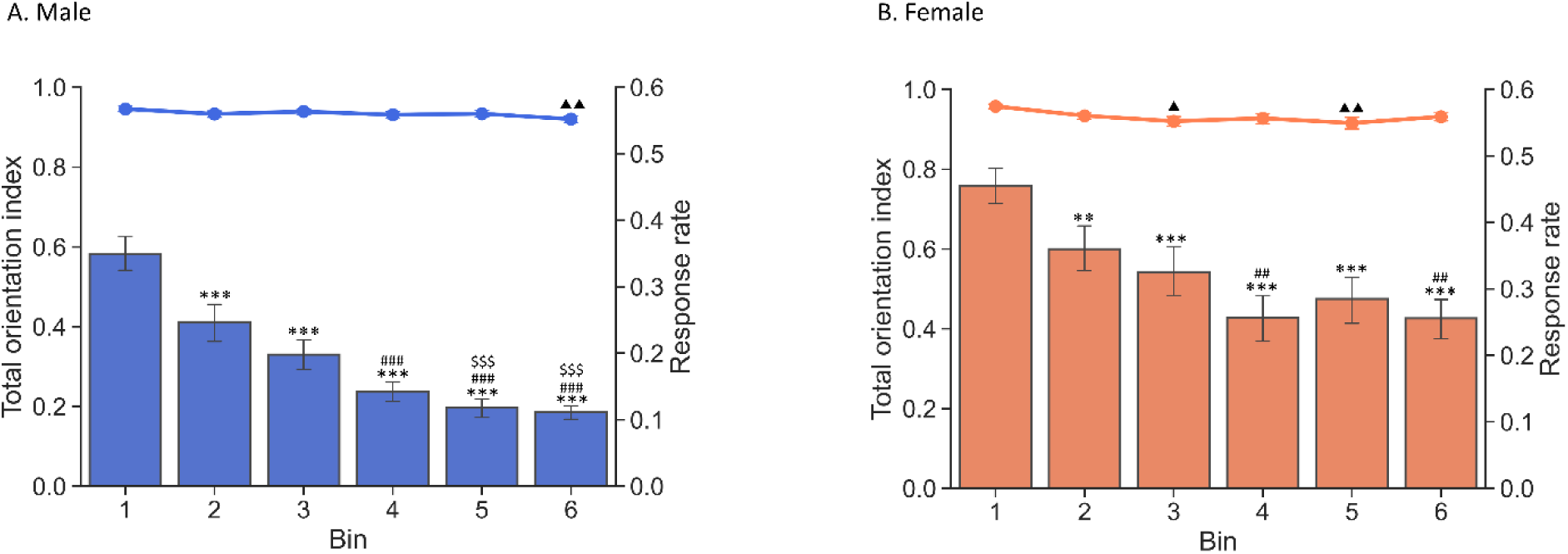
Task engagement in limited hold period is not affected by TOT. The line plot presents the total orientation index. The bar plot presents the response rate. The decreased response rate was not correlated with the change in orientation. Data are shown as mean ± SEM. n = 15 males and 7 females. A triangle (▴) in total orientation index curve indicates significant pos hoc results comparing to bin 1. An asterisk (**\***) in response rate bar indicates significant pos hoc results comparing to Bin1. A hash (#) in response rate bar indicates significant pos hoc results comparing to Bin2. A dollar sign (**$**) in response rate bar indicates significant pos hoc results comparing to Bin3. \*\**p*<0.01, \*\*\**p*<0.001, ##*p*<0.01, ###*p*<0.001, $$$*p*<0.001, ▴ *p*<0.05, ▴ ▴*p*<0.01.

In a comparison of the orientation index across response types, there was no effect of time (F(5, 460) = 1.0023, *p* = 0.4158), however, there was a main effect of response types on orientation index (F(3, 460)=130.6359, p< 0.0001), with significant pairwise differences between correct rejection, hit and miss (all *p* < 0.001), but not between hit and false alarm trials (*p* = 0.3798). There was also an interaction between time X response types (F(15, 460) = 1.9205, *p* = 0.0196) (Supplemental Figure 4A).

Similar to male mice, female mice showed a flat trend of total orientation index across time. There was a significant main effect of time bin (F(5, 50) = 3.7817, *p* = 0.0056). The pairwise comparisons showed significant difference between bin 1 vs bin 3 (*p* = 0.0152) and bin 1 vs bin 5 (*p* = 0.0040) (Figure 6B) with lower orientation indices during the later time bins. However, when combining the response rate results, a similar divergence to male mice was also present. The response rate started to decrease earlier than the orientation index, starting with Bin 2 (F(5,50) = 18.7840, *p* < 0.001; Bin1 vs Bin2, *p* = 0.0055; Bin1 vs Bin3, *p* = 0.0001; Bin1 vs Bin4, *p* < 0.0001; Bin1 vs Bin5, *p* < 0.0001; Bin1 vs Bin6, *p* < 0.0001).

The response type orientation index analysis of female mice also showed a main effect of response type (F(3, 229.04)=120.2993, p < 0.0001). There was no main effect of time bin and interaction shown (F(5, 229.04)=1.1346, p=0.3428; F(15, 229.04)=0.5567, p=0.9053, respectively) (Supplemental Figure 4).

### Amphetamine does not modulate physical orientation to the touchscreen

Because treatment with AMPH significantly improved performance in the rCPT, we further examined its impact on the orientation index in mice. Although a significant main effect of dose on the orientation index was observed in male mice (F(3, 293.63) = 5.7317, *p* = 0.0008) with the 0.3 mg/kg dose decreasing orientation compared to the 0.6 mg/kg (*p* = 0.0009) and 1 mg/kg (*p* = 0.0217) doses. Compared to the vehicle treatment group, none of the AMPH doses exhibited a significant effect on the orientation index (vehicle vs. 0.3 m/kg, *p* = 0.8440; vehicle vs. 0.6 mg/kg, *p* = 0.0984; vehicle vs. 1 mg/kg, *p* = 0.8795). Consistent with the orientation index results in the TOT probe, a small, but statistically significant decrease in orientation was noted over time (F(5, 292.19) = 3.4626, *p* = 0.0047), but only between Bin 1 and Bin 6 (*p* = 0.0013) (Figure 7A). The response rate decreased with time (F(5, 292.04) = 19.0334, p <0.0001), but AMPH did not affect the response rate (F(3, 293.30) = 0.6524, *p* = 0.5820),

**Figure 7.**
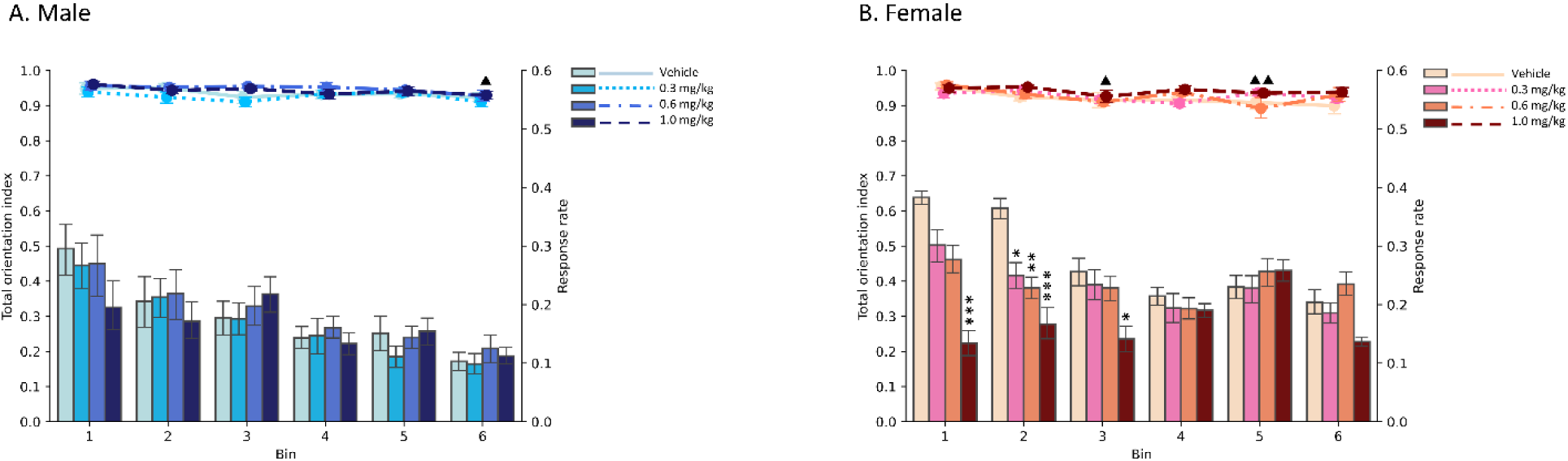
AMPH does not affect the orientation index during the limited hold period. The line plot presents the total orientation index. The bar plot presents the response rate. **A** The orientation index and response rate of male mice after amphetamine administration. AMPH did not alter the orientation index or response rate. **B** The orientation index and response rate of female mice after amphetamine administration. AMPH did not affect the orientation index in female mice. However, AMPH significantly decreased the response rate of female mice. Data are shown as mean ± SEM. n = 15 males and 7 females. A triangle (▴) in total orientation index curve indicates significant pos hoc results comparing to bin 1. An asterisk (**\***) in response rate bar indicates significant pos hoc results comparing to vehicle. \**p*<0.05, \*\**p*<0.01, \*\*\**p*<0.001, ^▴^*p*<0.05, ^▴ ▴^*p*<0.01.

Next, we evaluated the individual response types in male mice. Similar to the total orientation index results, there was no effect of AMPH treatment on orientation during hit or false alarm trials (F(3, 293.88) = 2.0177, *p* = 0.1115; F(3, 255)=0.9865, *p* = 0.3997, respectively). However, AMPH modulated orientation during misses and correct rejections (F(3, 294.18) = 4.8645, *p* = 0.0026; F(3, 294.11) = 3.9835, *p* = 0.0084, respectively), though the only significant differences were between 0.3mg/kg and 0.6mg/kg (in misses, *p* = 0.0025; in correct rejections, *p* = 0.0102, respectively) where orientation following 0.3 mg/kg treatment was lower. Notably, a small but significant decrease in the orientation index was observed only in correct rejections across time (F(5,292.22) = 3.4322, *p* = 0.0050), with significant decreases noted between Bin 1 and Bin 6 (*p* = 0.0109) and Bin 2 and Bin 6 (*p* = 0.0380) (Supplemental Figure 5).

In female mice, similar outcomes were observed in the orientation index under AMPH treatment, with no significant drug effect (F(3,137.04) = 2.2860, *p* =0.0815). Consistent with the previous TOT session results, orientation decreased over time (F(5, 137.04) = 3.0162, *p* = 0.0129) with significant differences between Bin 1 and Bin 3 and Bin 1 and Bin 5 (*p* = 0.0342; *p* = 0.0468, respectively) (Figure 7B). There were significant main effects of AMPH and time bin on response rates (F(3, 137.01) = 12.3878, *p* < 0.0001; F(5, 137.01) = 4.7873, *p* = 0.0005; respectively). The interaction between AMPH and time bin was also significant (F(15, 137.01) = 2.4779, *p* = 0.0030) (Figure 7B). The 1 mg/kg dose decreased the response rate in Bins 1, 2, and 3 compared to vehicle treatment (*p* < 0.001; *p* = 0.001; *p* = 0.0462; respectively). In Bin 2, the 0.3 and 0.6 mg/kg doses decreased the response rate compared to vehicle treatment (*p* = 0.0453; *p* = 0.0097, respectively).

When splitting the total orientation index into individual response types in female mice, results were consistent with those in male mice; no effect of AMPH was observed on hits or false alarms (F(3,140) = 0.7373, *p* = 0.5315; F(3,124) = 1.3267, *p* = 0.2688, respectively). AMPH treatment increased orientation during misses (F(3,137.02) = 5.4332, *p* = 0.0015), with the 1 mg/kg dose significantly increasing the orientation index compared to vehicle treatment (*p* = 0.0006). In contrast, AMPH did not affect orientation during correct rejections in female (F(3,137.05) = 1.6815, *p* = 0.1739) (Supplemental Figure 5B).

## Discussion

### The acquisition of rCPT training stage

The rCPT is a touchscreen-based measure of sustained attention, with strong face validity for human CPTs [11,26]. This study confirmed previous data from our group that both male and female C57BL/6J mice can reliably discriminate target and non-target stimuli and exhibit stable baseline performance using a truncated training procedure [15]. This training procedure is amenable to long-term drug screening studies with a single cohort of mice.

### Time-on-task effect on rCPT performance

By extending the session duration from 45 minutes to 90 minutes, the current study optimizes the rCPT for measurement of TOT-dependent decrements in performance in male mice. We observed a significant vigilance decrement, characterized by decreased sensitivity to the target throughout the session. Additionally, a shift from a liberal to a more conservative response bias was evident, with decreases in HR and FAR. These features parallel the findings in human literature related to TOT effects on performance [20]. A study utilizing a mouse 5C-CPT with extended sessions similarly demonstrated a decrease in the sensitivity index (non-parametric version of d’), but no change in the responsivity index (non-parametric version of c) [27]. This discrepancy may result from significant differences in the respective designs of the rCPT and 5C-CPT. For example, the rCPT requires both stimulus detection and discrimination while the 5C-CPT requires only stimulus detection. The 5C-CPT uses uniform light stimuli, while rCPT uses different images as targets and non-targets. Also, the 5C-CPT uses five possible stimulus locations, which engages spatially divided attention, while rCPTs only utilize a central stimulus location.

Notably, our results also suggest that male mice exhibit greater sensitivity to TOT-dependent performance decrements than females. The underlying reasons for this sex difference remain unclear, but are possibly related to sex differences in incentive salience or the behavioral policy employed in the task [28–30], so further studies are required to determine the causes of the sex difference.

### Amphetamine attenuates the TOT-induced vigilance decrement

As an FDA-approved treatment for ADHD, AMPH significantly improves performance in CPTs across species [14,31–36]. Here, we investigated whether AMPH could attenuate the vigilance decrement induced by TOT. Our findings demonstrate that in male mice, AMPH significantly improves sensitivity by reducing the FAR, while in female mice, despite significant decreases in the FAR, overall sensitivity remains unchanged. Similar improvements in d’ driven by decreased FAR were observed in a previous rCPT study [14].

The observed changes in the FAR in males and females and HR, response bias, and premature responses in females indicate that amphetamine shifts mice toward a more conservative response strategy. Female mice show a more dramatic change in response strategy, from liberal to conservative, compared to male mice. This may be due to inherent differences in baseline response strategies between sexes or sex differences in the pharmacokinetics or pharmacodynamics of AMPH.

### Changes in orientation to stimuli under TOT conditions

In this study, we integrated DeepLabCut-based behavioral tracking [23] and Visual Field Analysis [24] to accurately determine the orientation toward the touchscreen in the limited hold period of each trial across the TOT sessions. This approach allowed us to infer whether mice were physically engaged with the task. This neural network-based analysis platform offers unprecedented insights into performance during the rCPT and, potentially, other touchscreen-based behavioral tasks. To our knowledge, this is the first implementation of this type of analysis within the rCPT framework. Intriguingly, our findings indicate that both male and female mice maintained relatively stable physical engagement, measured as the total orientation index, within sessions, despite a small decline over time. The decline in the orientation index occurs much later in the session than the decline in response rate and d’ or the increase in c, suggesting that changes in performance are not driven by mice choosing to engage in competing behaviors during the session. Further analyses across different response types revealed no significant effect of time bins on physical engagement during any response type for either gender, reinforcing the fact that physical engagement is stable within rCPT sessions. This evidence supports the view that the changes in sensitivity and response bias represent a bona fide TOT-dependent vigilance decrement and are not due to physical disengagement from the task.

Cognitive psychology literature has extensively explored vigilance decrements related to TOT, proposing several theoretical models to explain this phenomenon, including resource depletion [37], mindlessness or mind-wandering [38–40], and a hybrid resource-control theory [41]. In the mindlessness theory, the vigilance decrement is due to boredom, or disinterest. In this case, reward and motivation can significantly affect task performance. A challenge in rodent sustained attention tasks is the potential decrease in motivation over time due to the availability of food rewards following trial completion, particularly in extended CPT versions designed to assess TOT effects. In our study, C57BL/6 male mice received approximately 2600μl (130 rewards of 20 µL each) of strawberry milk across the TOT session. Philips and colleagues tested C57BL/6J mice under a Fixed ratio 5 schedule with no trial limit. The results show mice using strawberry milk as a reinforcer complete an average of 150 FR5 trials, receiving about 3000 µL strawberry milk [42]. This suggests that 2600 µL does not produce complete satiety, however, we cannot rule out decreasing motivation over time as a cause of the TOT-dependent decrement in performance. However, given the consistent orientation index under TOT conditions, our data does not support a decrease in motivation or physical disengagement as the underlying causes for the vigilance decrement during the TOT-rCPT task. It is possible that satiety and the concomitant decrease in motivation for food produces “mind-wandering” that we are not able to accurately measure with the DLC-VFA platform.

### The effect of amphetamine on task engagement under TOT condition

Given that AMPH improves rCPT performance, we investigated whether AMPH impacts the orientation index under TOT conditions. Interestingly, AMPH did not alter the orientation index in either male or female mice, suggesting that AMPH does not modulate physical engagement. Notably, AMPH had no significant impact on the orientation index during most trial types compared to vehicle, except for the 1 mg/kg dose which increased the orientation index during misses in female mice. This result enriches our understanding of AMPH’s effects on female mice, particularly in light of previous observations where 1 mg/kg amphetamine increased the number of misses. Given that the orientation index during miss events did not decrease but rather increased, we can say that the increase in misses after amphetamine administration does not result from a lack of physical engagement. The increased orientation index during misses further corroborates the shift towards a more cautious approach, indicating that mice may more frequently observe the stimulus on the touchscreen before responding, rather than defaulting to a more liberal response strategy. The specific increase in orientation index for misses, but not other event types, could also be attributed to a potential ceiling effect, as the orientation values for other event types approach the maximum limit of the index.

In conclusion, by integrating markerless pose estimation (DLC) and visual field analysis (VFA), this study provides novel insight into the effects of TOT in the rCPT. We demonstrate that TOT induces a vigilance decrement not attributable to physical disengagement from the task and that AMPH attenuates this decrement by promoting a more conservative response pattern, significantly improving performance in male mice. Future studies will integrate these findings with techniques with the ability to provide information on electrophysiological activity during rCPT performance [43].

## Acknowledgements

We thank members of the Carr and Martinowich laboratories for helpful comments and suggestions. We also thank Aimee Ormond and Deveren Manley for assistance with animal care.

## Author Contributions

Conceptualization: YL, GVC

Formal analysis: YL

Investigation: YL, TvK, MM, BM, BSV

Writing-original draft: YL, GVC

Writing-review and editing: YL, TvK, MM, BM, BSV, JMB, JR, KM, GVC

Supervision: YL, KM, GVC

Project administration: KM, GVC

Funding acquisition: KM, GVC

## Supplemental Information

### Video recording and preprocessing

Videos were recorded using the original CCTV recording system attached to the touchscreen chambers. After cohort experiments were finished, all rCPT videos were exported from the CCTV unit for preprocessing. First, because the CCTV used variable frame rate (VFR) recording format, we converted the video clips to a consistent frame rate (CFR) format. When the CCTV unit recorded the video, each file had a size limitation of approximately 1GB. Once a file reached this limit, the recording would continue in a new file. As a result, there were multiple video files for each day. To facilitate the analysis, we first combined the video clips from each day into a single file. Subsequently, we trimmed the specific session videos from the combined file based on the timestamp, preparing them for further analysis using DeepLabCut. The ffmpeg package (Version 4.2.7) was used for preprocessing. The first frame of a session was defined as the first frame in which the reward tray was illuminated. Due to different touchscreen chamber configurations, the resolution of the video was either 960X480 (60FPS), 720X480 (60FPS) or 720X480 (30FPS).

### Pose estimation using DeepLabCut

For pose estimation, we used DeepLabCut (DLC; Version 2.2) <u>(Mathis et al. 2018; Nath</u> <u>et al. 2019)</u>. DLC was installed within the TensorFlow Docker Image with GPU support and executed in the Docker environment on the Ubuntu 20.04 platform. We used three different touchscreen configurations that each had different cameras and unique lighting conditions, so we trained three ResNet-50 based neural network models by using the labeled frames taken from each set. For the training dataset, 10-30 videos from each chamber set were used. 50 frames were taken from each video. Specifically, we labelled three body parts (Top of the head, left-side of the head and right-side of the head) in the DLC GUI interface. 95% of these labelled frames were used for training. The target evaluation result for each model was 1∼4 pixels for both training error and testing error (image size was 960X480 or 720X480). For each model, at least once model refinement was done. The outlier frames (frames with misidentified body parts) were picked out from the analyzed video by using the Jump and Manual algorithms See https://deeplabcut.github.io/DeepLabCut/docs/standardDeepLabCut_UserGuide.html for details. After relabeling body parts in outlier frames, these frames were merged into the training dataset and models were re-trained prior to re-analyzing the video.

### Visual Field Analysis (VFA)

Once pose estimation was completed in DLC, VFA was conducted (Mathilde Josserand et al., 2022). The VFA package was run using Spyder (Version 4.0.1, with Python 3.7.6, Qt 5.9.6 and PyQt 5.9.2) on Windows 10 platform.

After executing main coordinator module, the settings in the window were as follows: stimuli location > Left and/or right; stimulus fixed or moving > fiexed; group your frames by seconds > No; threshold for detecting outliers > 3; frontal angle for the animal > 20; lateral angle for the animal > 103.4.

Next, the program opened the first frame of each video in a pyplot interface. We marked four corners of chamber arena and specified the bottom and top border position of the touchscreen for each video. Prior to visual field analysis, two methods were applied to filter out outliers. In the first method, based on DeepLabCut predicted body part axis data in each frame, the following distances were calculated: top head to left ear; top head to right ear; left ear to right ear in each frame. Then, mean value and standard deviation of each distance was calculated based on all frames. Z score was then calculated by:

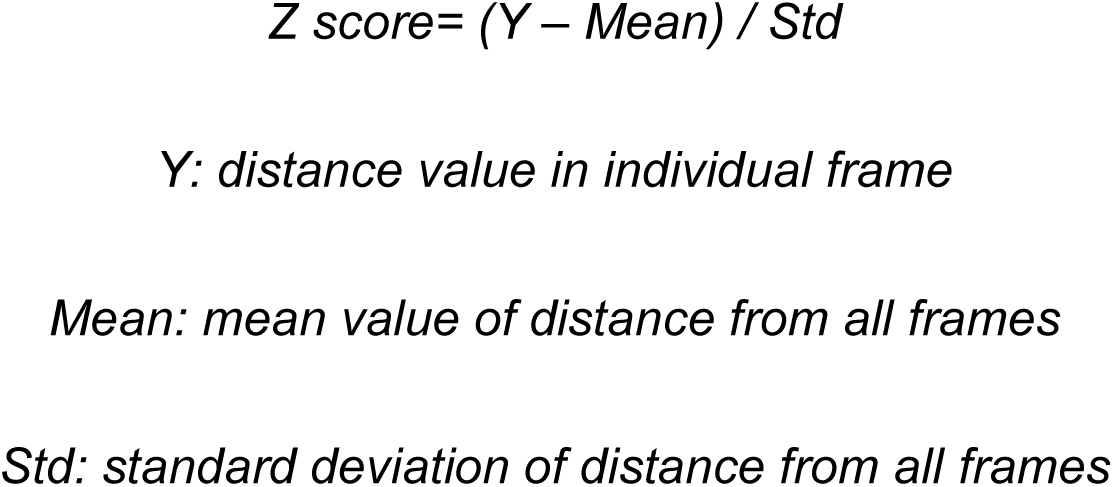

Any frame’s Z score > threshold indicated previously (set to 3) was excluded from analysis based on the recommendation from code developer (See https://github.com/mathjoss/VisualFieldsAnalysis/blob/master/modules/interface.py) The other method used the likelihood ratio reported by DeepLabCut results. The frame with likelihood lower than 0.9 in any body parts (top head, left ear and right ear) was excluded for further analysis. The program then processed the visual field analysis and exported the results into a .csv file. The following visual field were computed: frontal, lateral left, lateral right, blind, left all and right all. 0 ∼ 1 values were generated in each visual field to indicate the ratio of stimulus (or part of it) was in this visual field. If one frame showed that ≥ 90% of stimulus area (center area of touchscreen) was within blind visual field of mouse, we labeled this frame as blind frame, otherwise, we labeled this frame as oriented frame. Next, we identified the orientated trials based on the condition that at least one frame of video in the limited hold period was oriented frame. Further, we introduced a parameter named orientation index to present the animal orientation to the stimulus on the screen. The orientation index was calculated by using the number of orientated trials divided by total number of trials in each 15-minute time bin.

### Supplementary Figures

**Supplemental Figure 1.**
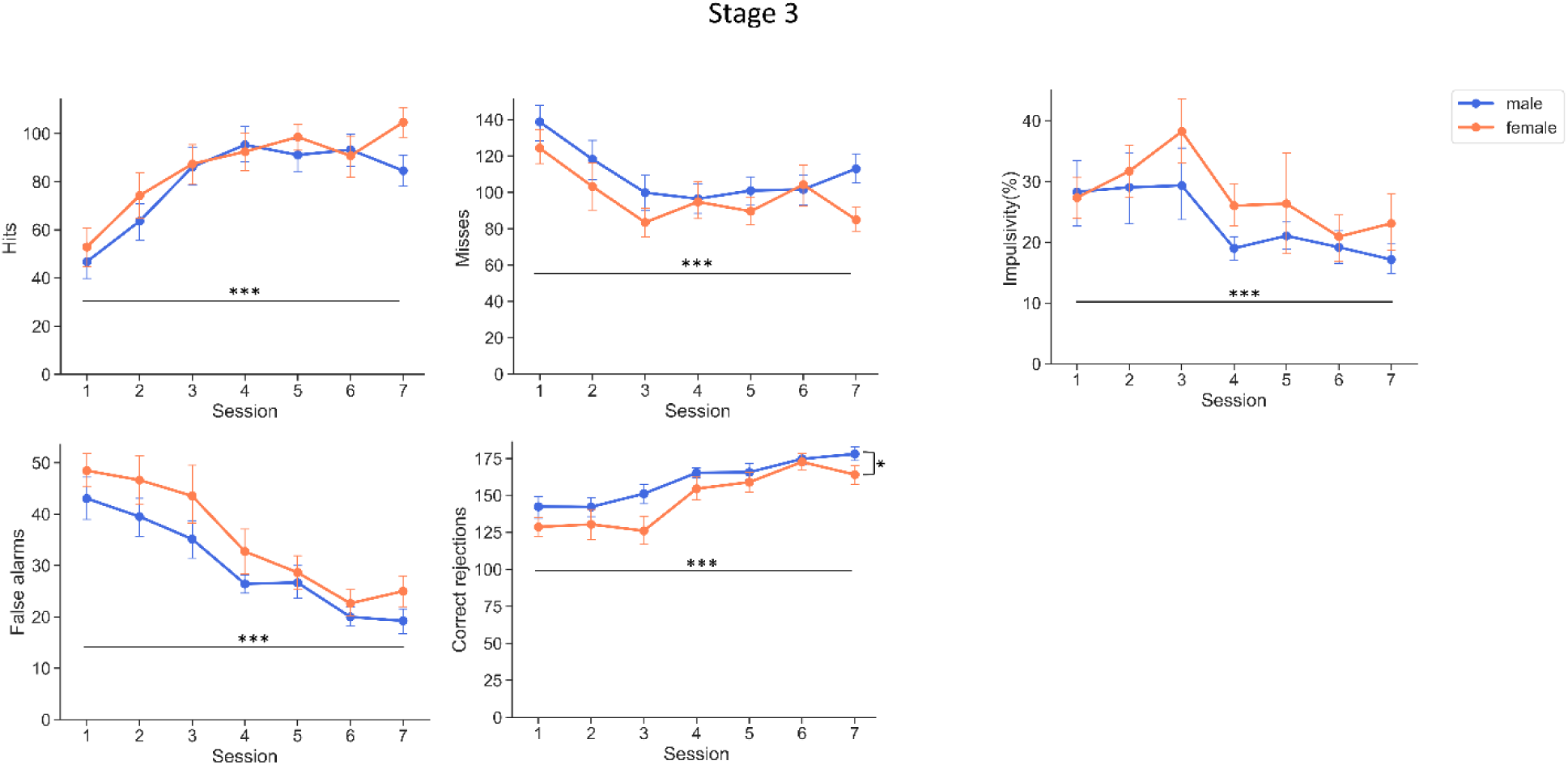
rCPT performance during Stage 3 training. Raw counts for each of the four possible rCPT responses and the impulsivity metric. Data are shown as mean ± SEM. n = 21 males and 11 females. An asterisk (*) indicates statistical significance of main effect of session or sex. *p<0.05, **p<0.01, ***p<0.001.

**Supplemental Figure 2.**
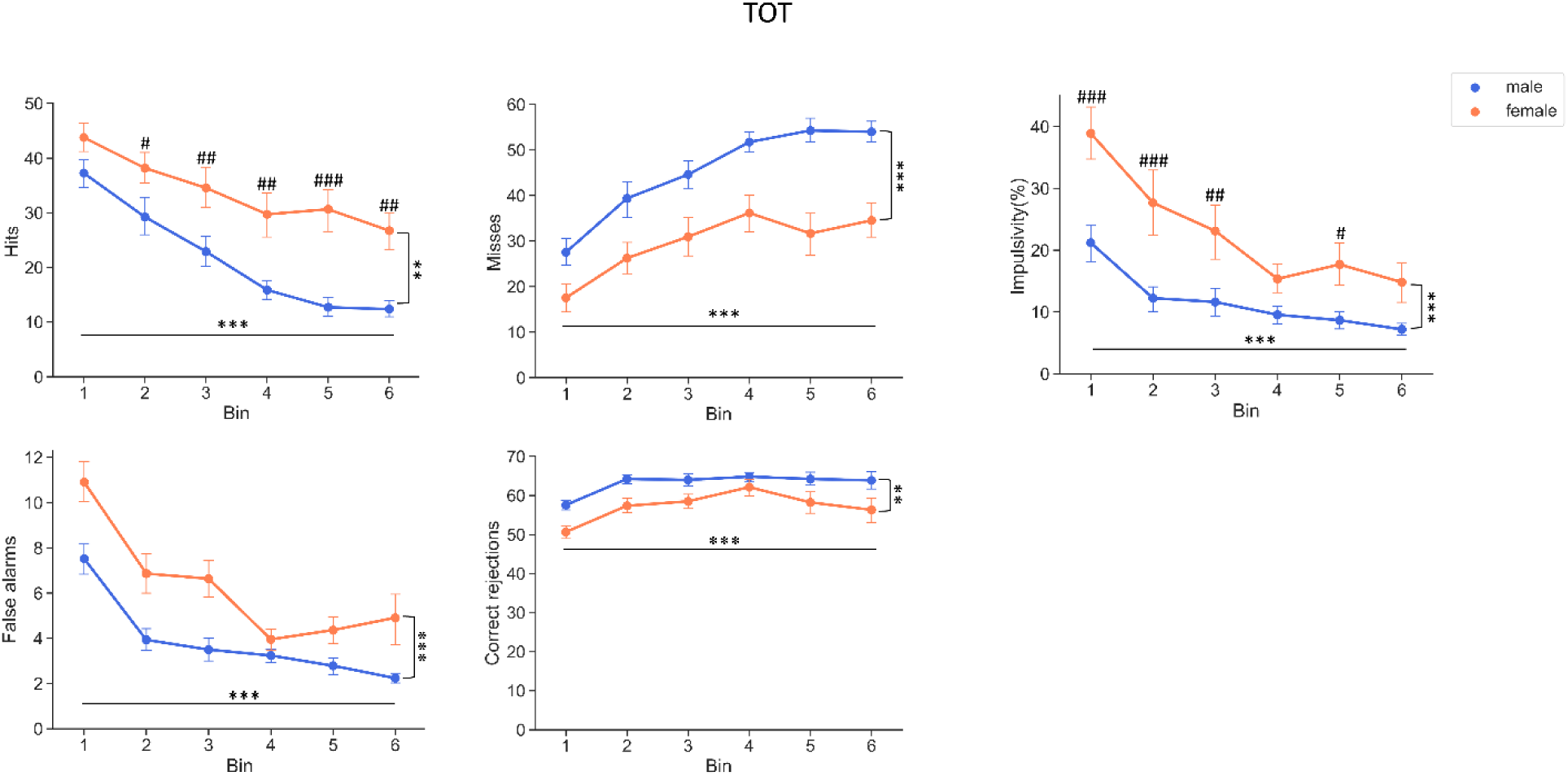
rCPT performance during TOT probe trials. TOT probe sessions were divided into six equal time bins of 15 minutes. Data are shown as mean ± SEM. n = 21 males and 11 females. An asterisk (*) indicates statistical significance of main effect of time bin or sex. A hash (#) indicates significant pos hoc results of time bin X sex interaction. *p<0.05, \*\**p*<0.01, \*\*\**p*<0.001, #*p*<0.05, ##*p*<0.01, ###*p*<0.001.

**Supplemental Figure 3.**
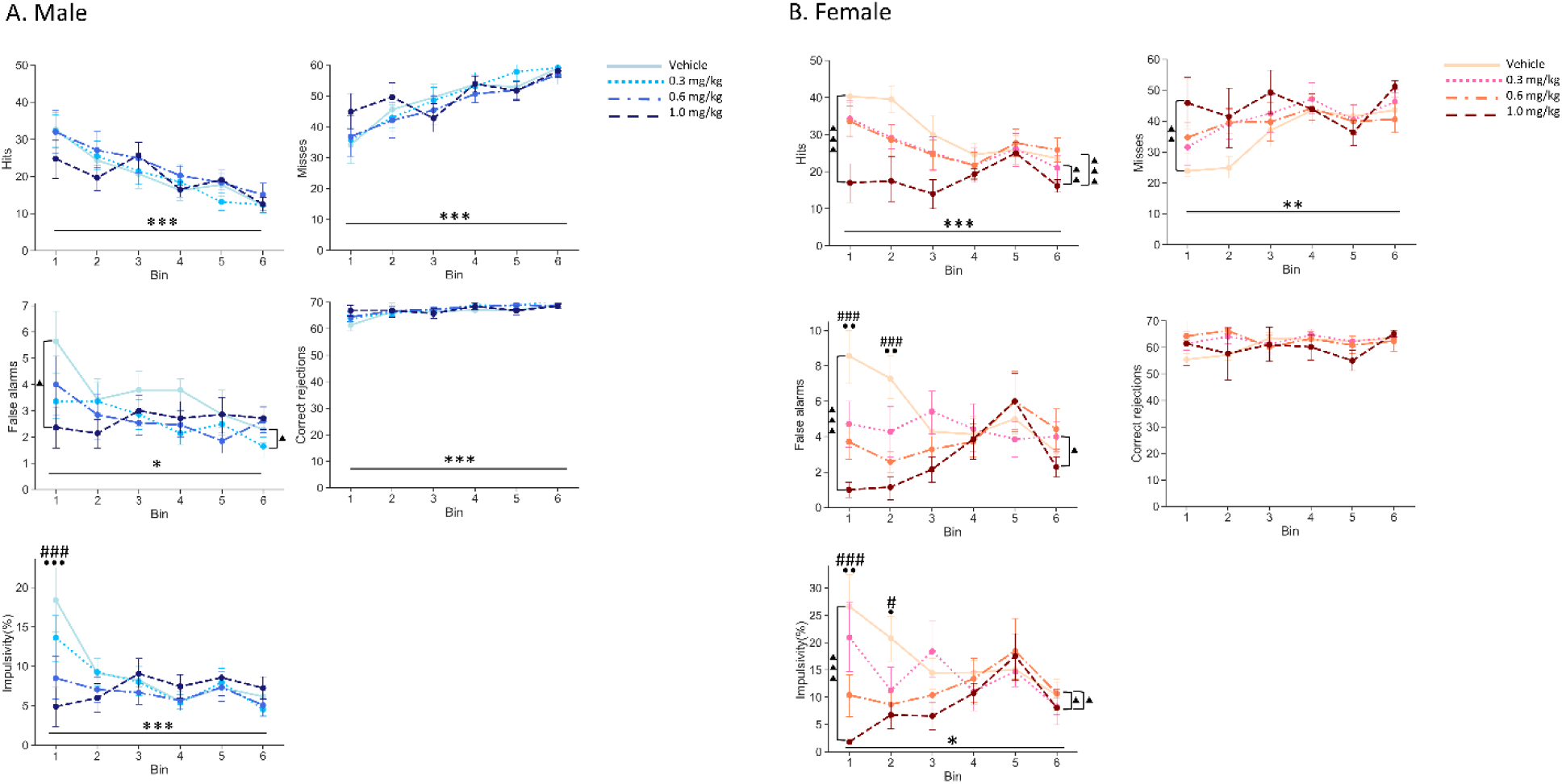
Effects of AMPH on rCPT performance. **A.** Male **B.** Female. An asterisk (*) indicates statistical significance of main effect or time bin or drug. A triangle (▴) indicates significant pos hoc result. A circle (•) indicates the significant effect of 0.6 mg/kg and a hash (#) indicates significant effect of 1 mg/kg after post hoc results of time bin X drug interaction. Data are shown as mean ± SEM. n = 15 males and 7 females. *p<0.05, \*\**p*<0.01, \*\*\**p*<0.001, • *p*<0.05, •• *p*<0.01, ^##^*p*<0.01, ^###^*p*<0.001.

**Supplemental Figure 4.**
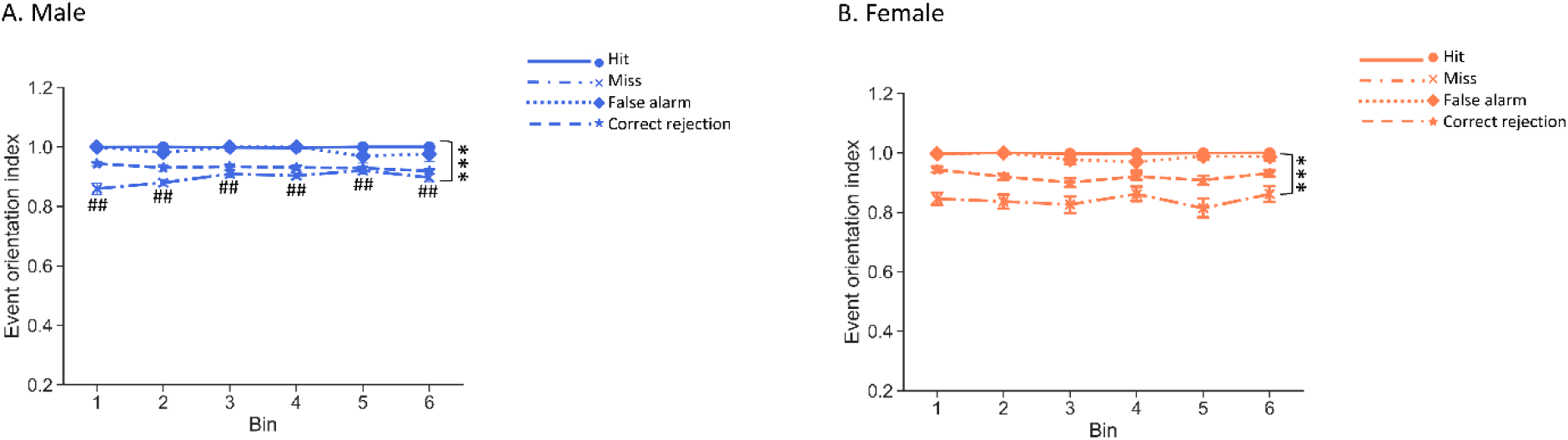
Task engagement during TOT probe sessions. **A.** Males **B.** Females Data are shown as mean ± SEM. n = 21 males and 11 females. An asterisk (*) indicates statistical significance of main effect of response type. A hash (#) indicates significant pos hoc results of time bin X response type interaction. \*\*\**p*<0.001, ##*p*<0.01.

**Supplemental Figure 5.**
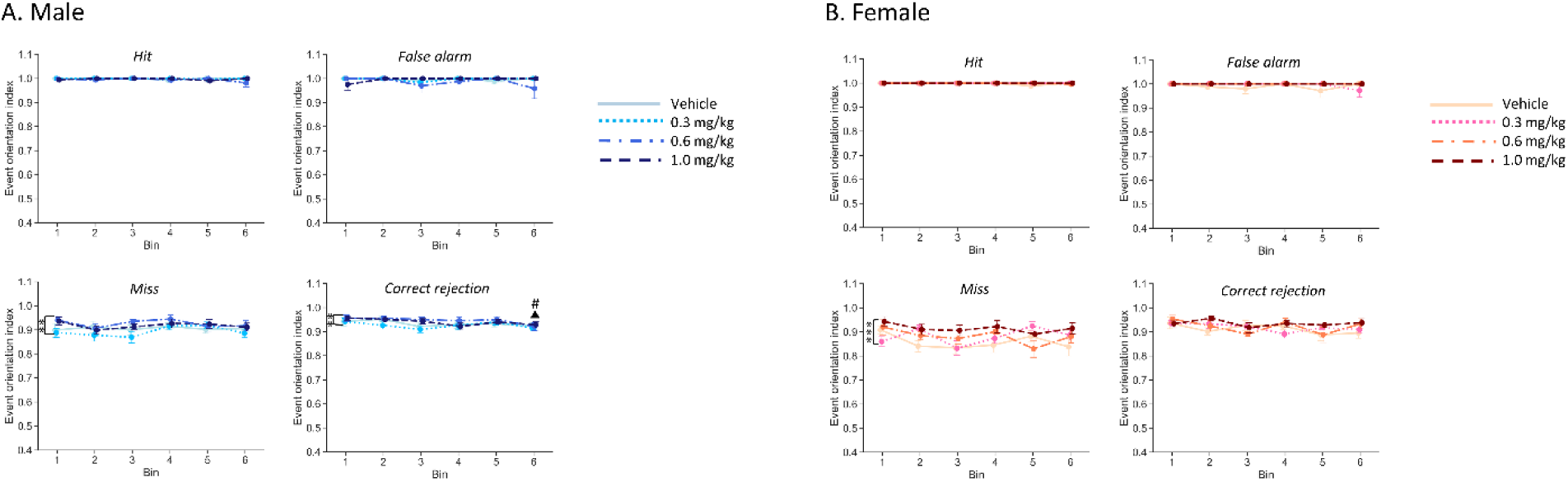
AMPH effects on the orientation indices during the limited hold period. **A.** Males **B.** Females. Data are shown as mean ± SEM. n = 15 males and 7 females. A main effect of time bin where there was a decrease in orientation during correct rejection trials between Bin 1 and Bin 6 (indicated by ▴); Bin2 and Bin 6 (indicated by #). B The orientation index of female mice after amphetamine administration. Main effect of drug was shown in orientation index of miss (indicated by*). \**p*<0.05, \*\**p*<0.01, \*\*\**p*<0.001, ^▴^p<0.05, ^#^p<0.05.

**Supplemental Table 1.**
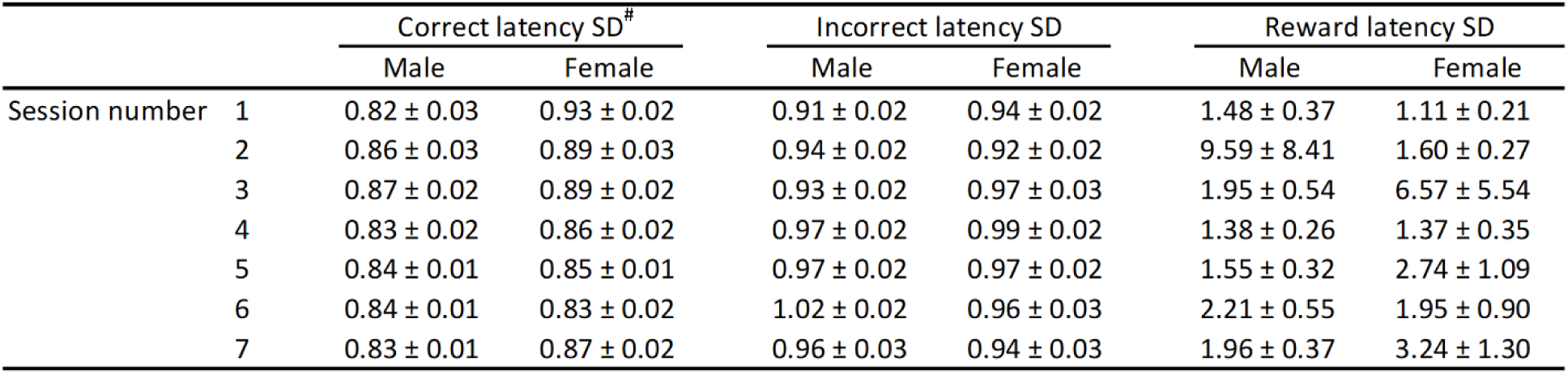

**Supplemental Table 2.**

|  |  | Correct latency SD |  | Incorrect latency SD |  | Reward latency SD |  |
| --- | --- | --- | --- | --- | --- | --- | --- |
|  |  | Male | Female | Male | Female | Male | Female |
| Time bin | 1 | 0.81 ± 0.02 | 0.80 ± 0.02 | 0.89 ± 0.05 | 0.89 ± 0.05 | 0.76 ± 0.11 | 0.40 ± 0.11 |
|  | 2 | 0.76 ± 0.02 | 0.84 ± 0.02 | 0.68 ± 0.08 | 0.81 ± 0.07 | 1.11 ± 0.38 | 0.65 ± 0.22 |
|  | 3 | 0.81 ± 0.02 | 0.84 ± 0.04 | 0.63 ± 0.08 | 0.86 ± 0.05 | 8.63 ± 6.47 | 0.82 ± 0.21 |
|  | 4 | 0.80 ± 0.04 | 0.78 ± 0.03 | 0.71 ± 0.07 | 0.75 ± 0.10 | 6.33 ± 2.75 | 9.28 ± 8.14 |
|  | 5 | 0.83 ± 0.03 | 0.80 ± 0.03 | 0.60 ± 0.08 | 0.75 ± 0.07 | 11.63 ± 6.30 | 10.3 ± 6.41 |
|  | 6 | 0.81 ± 0.03 | 0.84 ± 0.02 | 0.47 ± 0.07 | 0.73 ± 0.07 | 16.27 ± 8.39 | 6.98 ± 3.81 |

**Supplemental Table 3.**

| Male |  |  |  |  |  |  | Female |  |  |  |  |  |  |
| --- | --- | --- | --- | --- | --- | --- | --- | --- | --- | --- | --- | --- | --- |
| Dosage (mg/kg) | bin1 | bin2 | bin3 | bin4 | bin5 | bin6 | Dosage (mg/kg) | bin1 | bin2 | bin3 | bin4 | bin5 | bin6 |
| Correct latency SD |  |  |  |  |  |  | Correct latency SD |  |  |  |  |  |  |
| 0 | 0.78 ± 0.06 | 0.70 ± 0.07 | 0.84 ± 0.05 | 0.81 ± 0.04 | 0.84 ± 0.04 | 0.79 ± 0.06 | 0 | 0.85 ± 0.04 | 0.79 ± 0.01 | 0.84 ± 0.04 | 0.79 ± 0.04 | 0.80 ± 0.02 | 0.70 ± 0.07 |
| 0.3 | 0.80 ± 0.03 | 0.79 ± 0.03 | 0.80 ± 0.03 | 0.66 ± 0.07 | 0.74 ± 0.07 | 0.76 ± 0.04 | 0.3 | 0.79 ± 0.04 | 0.82 ± 0.03 | 0.85 ± 0.05 | 0.83 ± 0.03 | 0.81 ± 0.06 | 0.85 ± 0.05 |
| 0.6 | 0.71 ± 0.05 | 0.84 ± 0.03 | 0.82 ± 0.02 | 0.84 ± 0.04 | 0.72 ± 0.05 | 0.77 ± 0.05 | 0.6 | 0.81 ± 0.04 | 0.80 ± 0.03 | 0.88 ± 0.04 | 0.78 ± 0.04 | 0.85 ± 0.02 | 0.87 ± 0.02 |
| 1 | 0.70 ± 0.08 | 0.74 ± 0.07 | 0.74 ± 0.06 | 0.80 ± 0.04 | 0.78 ± 0.03 | 0.67 ± 0.06 | 1 | 0.63 ± 0.17 | 0.60 ± 0.16 | 0.63 ± 0.13 | 0.85 ± 0.02 | 0.81 ± 0.03 | 0.88 ± 0.04 |
| Incorrect latency SD |  |  |  |  |  |  | Incorrect latency SD |  |  |  |  |  |  |
| 0 | 0.75 ± 0.11 | 0.66 ± 0.13 | 0.66 ± 0.11 | 0.66 ± 0.10 | 0.41 ± 0.12 | 0.60 ± 0.12 | 0 | 0.80 ± 0.12 | 1.00 ± 0.07 | 0.80 ± 0.18 | 0.83 ± 0.14 | 0.93 ± 0.13 | 0.51 ± 0.24 |
| 0.3 | 0.63 ± 0.10 | 0.60 ± 0.11 | 0.48 ± 0.13 | 0.34 ± 0.11 | 0.60 ± 0.12 | 0.30 ± 0.11 | 0.3 | 0.56 ± 0.17 | 0.61 ± 0.23 | 0.67 ± 0.19 | 0.60 ± 0.19 | 0.86 ± 0.15 | 0.66 ± 0.13 |
| 0.6 | 0.59 ± 0.13 | 0.47 ± 0.14 | 0.46 ± 0.12 | 0.57 ± 0.12 | 0.38 ± 0.13 | 0.58 ± 0.13 | 0.6 | 0.48 ± 0.14 | 0.78 ± 0.17 | 0.70 ± 0.16 | 0.74 ± 0.07 | 0.66 ± 0.18 | 0.74 ± 0.19 |
| 1 | 0.29 ± 0.10<sup>▲</sup> | 0.24 ± 0.08<sup>▲</sup> | 0.64 ± 0.14 | 0.60 ± 0.13 | 0.55 ± 0.11 | 0.55 ± 0.13 | 1* | 0.37 ± 0.22 | 0.29 ± 0.21 | 0.52 ± 0.17 | 0.45 ± 0.15 | 0.92 ± 0.08 | 0.60 ± 0.16 |
| Reward latency SD |  |  |  |  |  |  | Reward latency SD |  |  |  |  |  |  |
| 0 | 0.31 ± 0.06 | 0.49 ± 0.15 | 1.11 ± 0.29 | 1.31 ± 0.32 | 1.44 ± 0.40 | 1.02 ± 0.27 | 0 | 0.66 ± 0.33 | 1.40 ± 0.55 | 2.63 ± 1.91 | 1.70 ± 0.49 | 1.54 ± 0.59 | 4.27 ± 2.72 |
| 0.3 | 1.29 ± 0.90 | 0.69 ± 0.20 | 1.24 ± 0.49 | 0.86 ± 0.23 | 0.88 ± 0.16 | 0.87 ± 0.17 | 0.3 | 0.55 ± 0.16 | 0.64 ± 0.14 | 2.14 ± 1.33 | 1.35 ± 0.61 | 3.08 ± 2.52 | 3.69 ± 1.64 |
| 0.6 | 0.94 ± 0.65 | 1.74 ± 1.28 | 1.14 ± 0.63 | 1.15 ± 0.30 | 1.53 ± 0.54 | 0.53 ± 0.12 | 0.6 | 0.68 ± 0.20 | 2.06 ± 1.26 | 9.16 ± 8.00 | 8.72 ± 8.03 | 0.90 ± 0.32 | 5.06 ± 4.59 |
| 1 | 1.44 ± 0.92 | 2.38 ± 1.74 | 1.56 ± 0.98 | 0.94 ± 0.29 | 1.24 ± 0.26 | 1.57 ± 0.39 | 1 | 0.28 ± 0.12 | 1.27 ± 0.93 | 13.3 ± 11.51 | 7.31 ± 5.65 | 8.72 ± 3.98 | 5.83 ± 2.41 |

